# Synergic action of Vitamin D_3_ and 17β-estradiol on behavioral impairments induced by unpredictable mild stress in the adult ovariectomized female rats

**DOI:** 10.1101/773739

**Authors:** Denis Baranenko, Vera Mironova, Julia Fedotova, Annemieke Johanna Maria van den Tol

## Abstract

The aim of the present study was study changes in antidepressant-like responses to Vitamin D_3_ (VD_3_) supplementation in three different doses (1.0, 2.5, and 5.0 mg/kg, s.c.) in long-term ovariectomized (OVX) rats treated with low dose of 17β-E_2_ (0.5 μg/rat, s.c.) during chronic unpredictable mild stress (CUMS). Sucrose preference (SPT), forced swimming (FST), and open-field (OFT) tests were administered to measure depression-like behavior. Brain-derived neurotrophic factor (BDNF), and serotonine (5-HT) and 5-hydroxyindoleacetic acid (5-HIAA) levels in the hippocampus were measured by ELISA, western blotting analysis, and by using high performance liquid chromatography (HPLC), respectively.

The findings demonstrated that VD_3_ (5.0 mg/kg, s.c.) in a combination with low dose of 17β-E_2_ increased sucrose consumption in the SPT and decreased depression-like in the FST of the long-term OVX rats exposed to CUMS. VD_3_ (5.0 mg/kg) resulted in significant elevated levels of hippocampal BDNF protein expression, as well as to the normalization of 5-HT and 5-HIAA levels in long-term OVX rats plus 17β-E_2_ compared to the OVX rats plus solvent or 17β-E_2_ submitted to CUMS. There were no significant effects of VD_3_ (1.0 and 2.5 mg/kg) treatment on both BDNF protein levels and 5-HT turnover in the hippocampus of the long-term OVX rats treated with 17β-E_2_ exposed to CUMS compared to the long-term OVX with CUMS rats plus solvent.

In conclusion, VD_3_ (5.0 mg/kg, s.c.) in a combination with low dose of 17β-E_2_ had a synergic antianhedonic- and antidepressant-like effects in the adult female rats following long-term ovariectomy submitted to CUMS. This is the first study in long-term OVX female rats showing beneficial effects of VD_3_ on depression-like behavior that are depended on the presence/absence of stressful factors.

## Introduction

Affective-related disorders, including depression are constantly rising, complicating the personal life style of people and increasing disqualification and hospital care [1,2]. Because of very high intensity of urbanization, our lifestyle and food choices have altered dramatically in the last twenty years [3,4]. Unfortunately these food modifications are associated to the marked scores of depression and other affective-related disorders in urbanized countries with high economic levels [5,6]. Moreover, in spite of numerous neuropharmacological schedules and psychological rehabilitation approaches for treatment of mood disorders, only moderate effects are registered in the treatment of affective-related disorders, and the possibility of recurrence is exalted especially in women [7,8].

It is generally recognized that women are two times more perceptive to different neuropsychiatric disorders than man [9,10]. Such sensitivity of women to mood disturbances are connected with female gonadal hormones fluctuations [8,10]. The findings of preclinical and clinical studies have shown that estrogen deficiency during perimenopause increase the susceptibility of women to affective-related disorders, including depression [8–10]. In this association, nutrients imbalance is considered as one of the critical causes enabling to the pathophysiological mechanisms for development of psychiatric disorders [5–7]. Application of additional nutritional interventions for treatment of mood deteriorations can be beneficial for both the prophylaxis and therapy of affective-related disorders [5–7,11].

VD deficiency has been noted in all people worldwide so it is postulated as a global problem [12–14]. Meaning that here is a growing interest of scholars and health practitioners worldwide in the investigators of the function of Vitamin D (VD) in human health and diseases, and especially in its pleiotropic outcomes [14,15].

VD is a fat-soluble prohormone that has skeletal and extra-skeletal functions [12,13]. The main sources of VD are sun exposure and diet. Two common forms of VD are VD_2_ (ergocalciferol) and VD_3_ (cholecalciferol) [15–17]. VD_2_ or VD_3_ must be activated to produce its effects under a multi-step process. VD may involve in the pathogenesis of affective-related disorders. Its active metabolic derivate 1.25 (OH) 2D exert both genomic and non-genomic actions [17,18]. This fact creates a neurobiological basis to propose the involvement of VD in the mechanisms of neuropsychiatric disorders [15–17]. VDR is a member of the nuclear hormone receptor super-family and acts as a ligand-inducible transcription factor as well as its non-genomic actions outside of the nucleus [17–19]. VDR has been detected in many brain regions, including the prefrontal cortex, hippocampus, cingulate gyri, thalamus, and hypothalamus [16–18]. The brain regions that contain VDR also demonstrated immunoreactivity for 1,25a-hydroxylase enzymes [15,16]. In this line, it can be assumed that 1,25-dihydroxyVD likely has humoral or neurohumoral activities in these brain structures [15–18].

On the other hand, there is a close interaction between the ovarian system and VD status, although detailed mechanisms of such an interaction are not fully identified and misinterpreted [20]. Low VD levels seem related to enhanced possibility for the induction of different gynaecological and obstetric disorders such as infertility, endometriosis, polycystic ovary syndrome, and breast or ovarian cancer [20]. Moreover, some clinical studies clearly show that mild depressive symptoms were more frequently registered in women with VD insufficiency, and especially, in women with VD deficiency [21–25]. Women in the perimenopausal period are more vulnerable to VD deficiency and its potential health consequences [23–26]. It is feasible that in perimenopausal and postmenopausal women, estrogen imbalance together with lower VD levels results in increased risk of negative outcomes for their health [21–24]. Moreover, our clinical studies have also demonstrated that young perimenopausal women (35-45 years) simultaneously showed high depression scores and profound decreased VD concentrations in the serum blood [27].

The pathogenesis of mood disorders is of multifactorial nature, including hormonal, genetics, inflammation, neurotrophins and/or monoamines imbalance [28–34]. Dysfunction of the hypothalamic-pituitary-adrenal axis (HPA) as hyperactivity is proposed to be one of the major provocative factors for initiation of affective-related disorders [28–30]. The other important factors for development of depression are decreased neurotrophins and/or serotonine (5-HT) levels in the brain [35–40]. The findings suggest that VD modulates production of neurotrophins, including brain-derived neurotrophic factor (BDNF), as well as the synthesis of amino acid tryptophan [13–18]. Low VD levels may accordingly induce low 5-HT contents [22,24].

Taking together all of the above-mentioned data, the affective-related disorders established in women with estrogen deficiency might be the result from multi-target alterations in the 5-HT system, VD_3_ levels, as well neurotrophins production. Our previous studies confirmed that VD_3_ (1.0 and 2.5 mg/kg, s.c.) plus low dose of 17β-estradiol (17β-E_2_) reduced depression-like behavior in the forced swimming test (FST) in adult non-stressed OVX rats following 12 weeks after surgery [41], suggesting that VD_3_ exhibits an antidepressant-like effects. Furthermore, we found that OVX rats after long-term ovariectomy demonstrated lower serum 25-OH-VD levels in the blood that are correlated with depression-like behavior [42,43]. However, it remains unclear if there is ameliorative antidepressant-like activity of VD_3_ on the chronic unpredictable mild stress model (CUMS) or whether the antidepressant-like action of VD_3_ involves BDNF and 5-HT system in the adult long-term OVX rats treated with 17β-E_2_ exposed to CUMS.

The aim of the present research was to study possible changes in the antidepressant-like responses to VD_3_ in long-term OVX rats treated with low dose of 17β-E_2_ subjected to CUMS. In this study, we used long-term estrogen deficiency caused by post-ovariectomy period for 3 months which is similar to research published previously [42,43]. This animal model is widely utilized in the preclinical behavioral research producing a menopausal-like state [44,45]. Sucrose preference test (SPT), forced swimming (FST), and open-field (OFT) tests, were performed to examine the depression-like state after VD_3_ treatment in the long-term OVX rats treated with low dose of 17β-E_2_ subjected to CUMS. Serum corticosterone (CS) levels, hippocampal 5-HT/5-HIAA and BDNF contents were examined to identify the possible mechanisms of VD_3_ effects on the behavioral profile in long-term OVX rats treated with 17β-E_2_ subjected to CUMS.

## Materials and methods

### Animals

Fifty-six female rats of the Wistar line at age 3 months (weighing 210 ± 20 g) were purchased from the Animal Rat Center of the Rappolovo Laboratory Animal Factory (St. Petersburg, Russia). All females were maintained under standard animal vivarium conditions with a constant room temperature (22 ± 1 °C), relative humidity (50 ± 10%), and a 12 h light/dark cycle (light from 07:00 to 19:00) with typical food for rodents and water ad libitum. All rats could habituate to the novel environment for 1 week prior to their use in this research. All stress manipulations were performed to minimize any pain and undesirable experiences in the experimental animals. The entire research procedure was approved by the Animal Care Committee of the I.P. Pavlov Institute of Physiology (Protocol No.: 1095/1/25.06.2012) and carried out in compliance with the National Institute of Health guidelines for laboratory animals.

### Ovariectomy

To modulate the hormonal state of the females Wistar rats a bilateral removal of the ovaries was conducted. It might be important to note that this hormonal state is considered to be similar to the menopausal period in women [44,45]. As part of this procedure a narcosis was performed by means of an intraperitoneal administration of xylazine 10 mg/kg and ketamine 70 mg/kg. This was followed by the removal of both ovaries. This procedure was performed by using two standard cuts in a lateral position. After this procedure the muscles and skin incision were restored by surgical staples. The efficacy of this surgery was validated by a routine vaginal inspection and inspection of serum estradiol levels. For the sham operation, the same procedure was followed but without the amputation of the ovaries. This entire procedure was performed in line with previously published research [42,43].

Following the ovariectomy or sham operation, the OVX rats were placed in their cage and were allowed to recover for a period of 12 weeks while having continues access to food and water. Following this time, each rat was randomly assigned to an experimental group for the chronic stress procedure, except for the sham-operated (SHAM), non-stressed, control rats.

### CUMS model

To induce clinical depression in the rats in the experimental conditions the chronic unpredictable mild stress (CUMS) procedure [46] was followed with some small alterations [47,48]. The CUMS procedure includes exposure to repeated unpredictable stressors which follow a four weeks protocol which is meant to appear random and unpredictable to the animals [49]. Animals in CUMS groups were single cage bred and subjected to different types of stressors which varied from day to day to make the stress procedure unpredictable. The list of stressors are: water deprivation (24 h), food deprivation (24 h), wet bedding (24 h), tilted cage at 45° (24 h), unpredictable shocks (15 mA, one shock/20 s, 10 s duration, 20 min), a reversal of the light/dark cycle (24 h), swimming in cold water (4 °C, 5 min), swimming in warm water (45°C, 5 min), and clipped tail (1 min, 1cm from the end of the tail). Each stress trigger was performed approximately 3 times during procedure protocol. Rats exposed to one of these stress triggers per day. All of the stressors were supplied individually and continuously for day and night. In order to avoid habituation the same stressor was not arranged consecutively for 2 days, which would guarantee the animal will not predict the occurrence of any stress trigger. CUMS protocol was similar every time to maintain reproducibility. During the stress process, rat was moved into experimental room to accept the stressor and returned its single cage after performing the stress procedure. The stress procedure did not involve any food or water deprivation in the present study.

The control, sham-operated female rats were housed in a separate cage (and room) without any contact with the stressed groups of animals. These rats were generally undisturbed and well maintained (provided with food and routine cage cleaning). All rats were weighed before and after the CUMS period (Fig. 1). Furthermore, all experimental groups of rats were supervised daily by the veterinary care staff during subroutine maintenance. The rats were examined in the special animal room using a noninvasive observational assessment procedure that yielded information regarding the health state of each animal. The assessment consisted of several measurements: body condition, appearance, breathing, hydration status, posture, mobility, muscle tone, and the presence of defects in the bones, genitals and abdomen. There exist no rats damaged or unhealthy during the experimental protocol. Timeline of this experimental study is presented on Figure 1.

**Fig 1.**
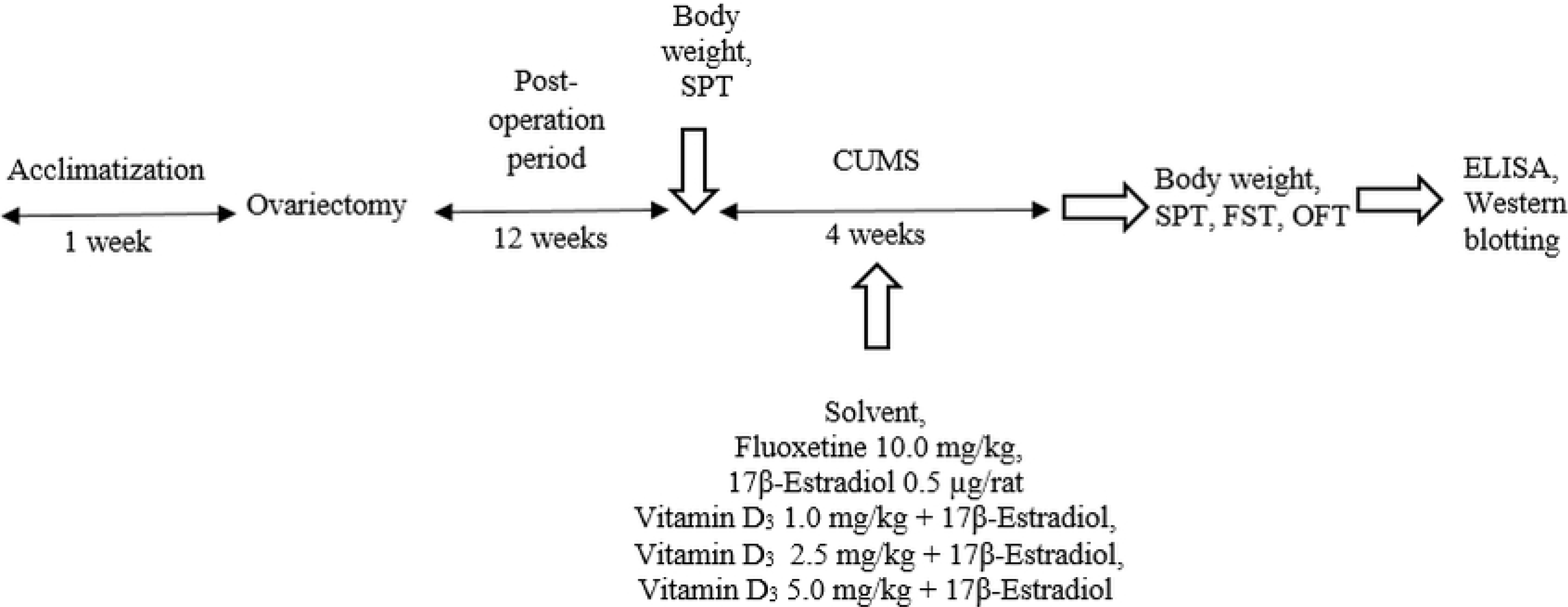
Timeline of chronic treatment. Female Wistar rats were divided into 6 groups – non-CUMS SHAM rats treated with solvent (control), SHAM rats submitted to CUMS treated with solvent, long-term OVX rats exposed to CUMS given with solvent, fluoxetine as positive control (10.0 mg/kg/day), 17β-E_2_ (0.5 μg/rat/day, s.c.) or VD_3_ (1.0, 2.5, 5.0 mg/kg/day, s.c.) in a combination with low dose of 17β-E_2_. All rats were weighed before and after the CUMS period.

### Drugs

The drugs used in this research were 17β-E_2_, fluoxetine hydrochloride, and VD_3_ as cholecalciferol. All drugs were obtained through Sigma Chemical Co. (St. Louis, MO, USA) and injected subcutaneously (0.1 ml/rat) for the 4 weeks during the CUMS procedure – 30 min before the daily stressor action – and throughout the period of the behavioral tests. Behavioral measurements were conducted 60 min after the final drug administration. Sterile sesame oil was used for the preparation of estrogen. Vitamin D_3_ was dissolved in a solution of 95% ethanol, and then aliquoted and kept at −80°C. Fluoxetine hydrochloride was dissolved in sterile physiological saline. Cholecalciferol (which was used for injection into the females in the experimental conditions) was diluted in sterile water, resulting in a solvent of VD_3_ with 2% of ethanol.

### Groups of animals

All animals were randomly assigned to the eight experimental groups (n = 7 in each): SHAM rats without the CUMS model treated with saline (control), SHAM rats submitted to CUMS treated with oil solvent, long-term OVX rats exposed to CUMS given with oil solvent, fluoxetine as positive control (10.0 mg/kg/day), 17β-E_2_ (0.5 μg/rat/day sc) or VD_3_ (1.0, 2.5, 5.0 mg/kg/day) in combination with 17β-E_2_. In our preliminary studies, there were no significant differences between SHAM/OVX rats treated with physiological saline as solvent for fluoxetine, and SHAM/OVX females treated with sterile water with 2% ethanol as solvent for VD_3_ or SHAM/OVX females treated with sesame oil as solvent for 17β-E_2_ in behavioral trials (data are not shown). Since, we did not find any differences between these experimental groups, the sesame oil as solvent for SHAM/OVX females was used in the present work. The doses of VD_3_ were based on our previous studies on the behavioral effects of VD_3_ on depression-like behavior of non-stressed long-term OVX female rats [42]. The dose of fluoxetine was utilized according to earlier experimental data [50]. Several studies have demonstrated that the administration of fluoxetine decreases depressive-like behavior in rodents [50,51]. All behavioral measurements were made 60 min after the last drug administration. To minimize animal suffering, all groups of rats were euthanized by pentobarbital overdose after all behavioral trials.

### Sucrose preference test

Before and after the initiation of the 4 weeks CUMS procedures, the experimental rats underewent the sucrose preference test (SPT) [52,53]. This test is set up as follows: Following a training trial, the rats are subjected to a 24 h deprivation of food and water. On the next day, the rats have one hour access to one bottle with 200 ml of water and a simalar amound of sucrose solution. The experimenter measures the percentage of the consumed sucrose solution and water volumes as a measure of sucrose preference by calculating the value of the sucrose preference among all (sucrose plus water in mL) liquid consumption:

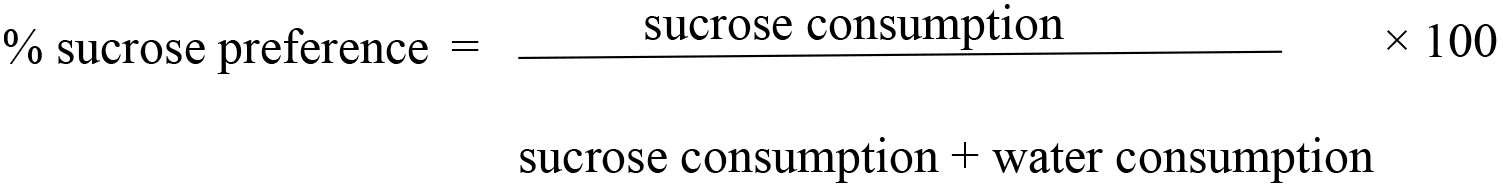

### Forced swimming test

For testing of modifications of depression-like behavior, all groups of rats were submitted to the standard forced swimming test (FST) as described in earlier work (FST) [42]. The 3 cylinders (60 cm tall and diameter 20 cm) were filled with 23 −25°C water up to a 30-cm depth. On the first day, the rats were pre-tested during 15 min in cylinders. Then, the rats were dried with papers and placed at their home cages till the next day. On the second day (testing trial), OVX females with CUMS were examined into the apparatus for 5 min. The following parameters were registered: 1) immobility time (floating in the water with only movements necessary to keep the head above water); 2) swimming time (active swimming movements around glass cylinder); 3) climbing time (active movements with forepaws directed towards the walls). For recording of these values, a video camera was installed above the apparatus.

### Open field test

The measurements of the behavioral activity in the OFT were carried out as in previously published research [42,43]. The rats were set in the center square of the OFT and tested for 5 min. Motor activity and rearing and grooming behavior were recorded for 300 s in the OFT apparatus using a video camera and the equipment was cleaned in-between sessions.

### Biochemical assay

After all the behavioral testing, all rats underwent a narcosis, and approximately 5 ml samples of blood were drawn from the animals to be centrifuged at 4000 g for 15 min at 4 °C. While doing so the hippocampi of rats in the experimental group were dissected to be homogenized in cold lysis extraction buffer (0.2% sodium deoxycholate, 0.5% Triton X-100, 1% NP-40, 50 mM Tris–HCl pH 7.4, 1 mM phenylmethylsulfonyl fluoride, 1 mM N-ethyl-maleimide, and 2.5 mM phenantroline) [54]. After that, the hippocampal samples with the cold lysis buffer were sonicated for 15 s. Then, the hippocampi were centrifuged at 12,000 g for 15 min at 4 °C. The Bradford method was used for the normalization of hippocampal supernatants to the total protein [55]. The serum samples and hippocampal protein normalized supernatants were stored at −80 °C until the ELISA assays. The serum samples were used for the measurement of the 25-hydroxyVD_3_ (25-OH-VD_3_), and corticosterone (CS) levels using a commercially available rat ELISA kit (Cusabio Biotech Co., Ltd, Wuhan, P.R. China) according to the manufacturer’s instructions. The sensitivity and detection range of the 25-OH-VD_3_ rat ELISA kits were 5.0 μg/l and 20–100 μg/l, respectively. The sensitivity and detection range of the corticosterone rat ELISA kits were 0.1 ng/ml and 0.2–40 ng/ml, respectively.

Hippocampal homogenates were used for the detection of the BDNF level by the rat ELISA kits (Cusabio Biotech Co., Ltd, Wuhan, P.R. China) according to the manufacturer’s instructions. Briefly, 100 *μ*L of hippocampal sample or standard was added to each well and incubated for 120 min at 37.0 ∘C. Then, 100 *μ*L of anti-BNDF antibodies were added to each different well and incubated for 60 min at 37.0 ∘C. After 3 times of washing, 100 *μ*L of HRP–avidin working solution was added to each well and incubated for 60 min at 37.0 ∘C. Again, after 5 times of washing, 90 *μ*L of tetramethylbenzidine solution was given to each different well and incubated for 15–30 min at 37.0 ∘C. Then, 50 *μ*L of stop solution was added to each well to terminate the color reaction. The BDNF levels were measured using a MC Thermo Fisher Scientific reader (Thermo Fisher Scientific Inc., Finland) with an absorbance of 450 nm. The standard curve was used for the calculation of the relationship between the optical density and the BDNF levels. The BDNF content is presented as pg/mg of tissue. The sensitivity and detection range of the BDNF rat ELISA kits were 0.078 ng/ml and 0.312–20 ng/ml, respectively. The assay exhibited no significant cross-reactivity with other neurotrophic factors. All samples were duplicated for the assay.

### HPLC detection of 5-HT and 5-HIIA levels in hippocampus

For the groups of OVX rats and OVX plus 17β-E_2_ rats subjected to CUMS the measurements of serotonin (5-HT) and its metabolite 5-hydroxyindoleacetic acid (5-HIAA) levels in the hippocampus were investigated in order to measure the 5-HT neurotransmission of the hippocampus as a response to the supplementation of VD_3_. The measurements of 5-HT/5-HIAA levels in the hippocampus were conducted with the use of high-performance liquid chromatography and electrochemical detection (HPLC) as in previously published research [56].The hippocampi were dissected on dry ice (similarly to for the ELISA) to obtain BDNF levels, samples were stored and weighed at −80 ◦C until during the entire examination. All hippocampal tissue was homogenized in a 0.1 mol/L solution of HClO_4_ which contained 0.02% Na_2_S_2_O_2_ (15 μL of solution for each milligram of tissue) and dihydroxybenzylamine (DHBA, 146.5 ng/mL, internal standard). These homogenates were centrifuged for a period of 40 min at 11.000 g at 4 °C. A Shimadzu LC-10AD (Kyoto, Japan) isocratic system was used with a Spheri-5 RP-18 5 μm column (220 × 4.6 mm) and a 20 μL injection loop, by electrochemical detection at 0.75 V and a mobile phase composed of 0.06% heptane sulphonic acid and phosphate/citrate pH 2.64, 0.02 mol/L, 0.12 mmol/L ethylene diamine tetraacetic acid, containing 10% methanol, at a flow rate of 1 mL/min. 5-HT and 5-HIAA levels were expressed as mean ± SD ng/mg tissue. Prior to the examination of the hippocampal tissue, a solution was prepared containing 2 mL of sodium tetraborate 0.1 mol/Land 1 mL of stock solution. The precolumn derivatization was finalized by reacting 100 μL of this solution with 50 μL of sample for 2 min before each injection (Donzanti, Yamamoto, 1988). The mobile phase entailed a sodium phosphate 0.05 mol/L (pH 5.95) with methanol 11.5%. The flow rate of the HPLC system was 3.5 mL/min and a detector was utilized with an emission of 460 nm and excitation of 348 nm. Standards of 5-HT and 5-HIIA were employed and to settle there were no extraneous peaks the retention time was validated for each substance.

### Western blotting analysis

Hippocampal tissues were homogenized in a cold lysis buffer containing a protease inhibitor cocktail (Sigma-Aldrich, USA) for 1 h and centrifuged at 12,000 g at 4 °C for 20 min. The protein content was evaluated by a Bio-Rad protein detector (Bio-Rad, USA), and 100 *μ*g of total protein from each sample was denatured with a buffer (6.205 mM Tris–HCl, 10% glycerol, 2% SDS, 0.01% bromophenol blue, and 50 mM 2ME) at 95 °C for 5 min. The denatured proteins were separated on a SDS page (10% sodium dodecyl sulfate polyacrylamide gel) and forwarded to a nitrocellulose membrane (Amersham Biotech, USA). After that, the membranes were probed with anti-BDNF (1:1000, Santa Cruz) and β-actin (1:1000; Sigma-Aldrich, USA) monoclonal antibodies for 2 h and secondary antirabbit antibodies (1:5000; Santa Cruz, USA) conjugated to horseradish peroxidase for BDNF for 1 h. Bands were detected by 5-bromo-4-chloro-3-indolyl phosphate with a nitro blue tetrazolium kit (Abcam, China) as a chemiluminescent substrate. Signals were measured by an image analysis system (UVIdoc, Houston, TX, USA).

### Statistical analysis

All experimental data are expressed as the mean ± standard deviation of the mean. The treatment effects were determined with a one-way ANOVA followed by an LSD *post hoc* test using the Statistics Package for SPSS, version 16.0 (SPSS Inc., USA). A value of P < 0.05 was considered statistically significant.

## Results and discussion

### VD_3_ alters the body weight in the long-term OVX rats treated with 17β-E_2_ exposed to CUMS

The body weights of long-term OVX rats subjected to CUMS and treated with 17β-E_2_ in a combination with all investigated doses of VD_3_ are presented in Figure 2. There was no difference in the initial body weight in all the experimental groups. Following 4 weeks, the body weight of SHAM rats with CUMS was significantly decreased compared to the control, non-CUMS SHAM group (F(1,34) = 72.66, P < 0.001; Fig. 2). The body weight of long-term OVX rats with CUMS was significantly decreased compared to the non-CUMS/CUMS SHAM groups (P < 0.001; Fig. 2). Administration of 17β-E_2_ did not statistically enhance body weight of long-term OVX rats with CUMS compared to the non-CUMS control, CUMS OVX/SHAM groups (P > 0.001; Fig. 2). However, there was a tendency to increase the body weight of long-term OVX rats with CUMS compared to the OVX rats plus CUMS given with solvent. Supplementation with VD_3_ (5.0 mg/kg) plus 17β-E_2_ significantly prevented the reduction of the body weight of long-term OVX rats with CUMS (P < 0.001; Fig. 2) compared to the OVX plus solvent or 17β-E_2_ /SHAM rats exposed to CUMS (P < 0.001; Fig. 2). This effect of co-administration of VD_3_ (5.0 mg/kg) plus 17β-E_2_ was similar to the effect of the reference drug fluoxetine (10.0 mg/kg) in long-term OVX rats with CUMS. VD_3_ at doses of 1.0 or 2.5 mg/kg in a combination with 17β-E_2_ failed to modify the body weight of long-term OVX rats with CUMS compared to the non-CUMS control, OVX/SHAM groups exposed to CUMS (F(1,34) = 0.28, P > 0.001; Fig. 2), which is consistent with the results in the SPT, FST, and OFT.

**Figure 2.**
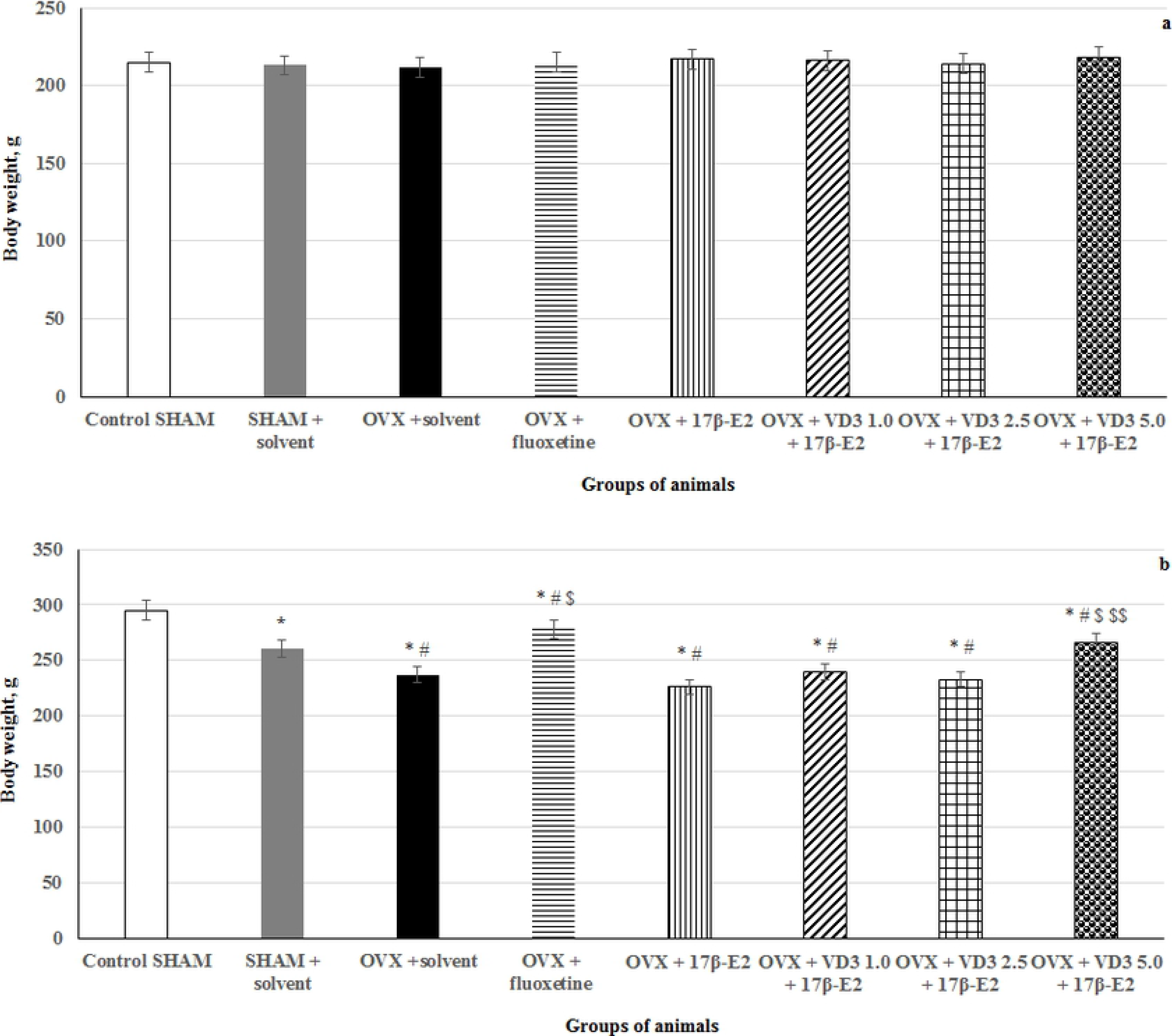
VD_3_ corrects the body weight in the long-term OVX rats treated with 17β-E_2_ exposed to CUMS. a – prior to CUMS, b – after CUMS. * – P < 0.05 versus the control group, # – P < 0.05 versus to the SHAM group with CUMS, $ – P < 0.05 versus to the OVX group with CUMS, $$– P < 0.05 versus to the OVX group with CUMS treated with 17β-E_2_. The data are presented as mean ± SD; n = 7 in each group.

### VD_3_ increases sucrose preference in the long-term OVX rats treated with 17β-E_2_ exposed to CUMS

Before the CUMS protocol, there was no significant difference among the experimental groups in the SPT (Fig. 3). Following 28 days of the CUMS trials, the SHAM rats exhibited a decrease in sucrose preference when compared to the control non-CUMS SHAM group (P<0.05). The sucrose preference in long-term OVX rats were significantly reduced compared to the non-CUMS/CUMS SHAM rats (F(1,34) = 56.14, P < 0.05; Fig. 3). Low dose of 17β-E_2_ increased sucrose preference in long-term OVX rats with CUMS compared to the OVX group with CUMS plus solvent (P > 0.05; Fig. 3). Treatment with VD_3_ at dose of 5.0 mg/kg plus 17β-E_2_, as well as fluoxetine, markedly increased sucrose consumption in the long-term OVX rats exposed to CUMS when compared to the OVX plus solvent or 17β-E_2_ /SHAM rats submitted to the CUMS (P < 0.05; Fig. 3). There was no signifcant differences among the groups of long-term OVX/SHAM females subjected to CUMS and the OVX rats with CUMS administered by VD_3_ at another two doses plus 17β-E_2_ (F(1,34) = 1.16, P > 0.05; Fig. 3).

**Fig. 3.**
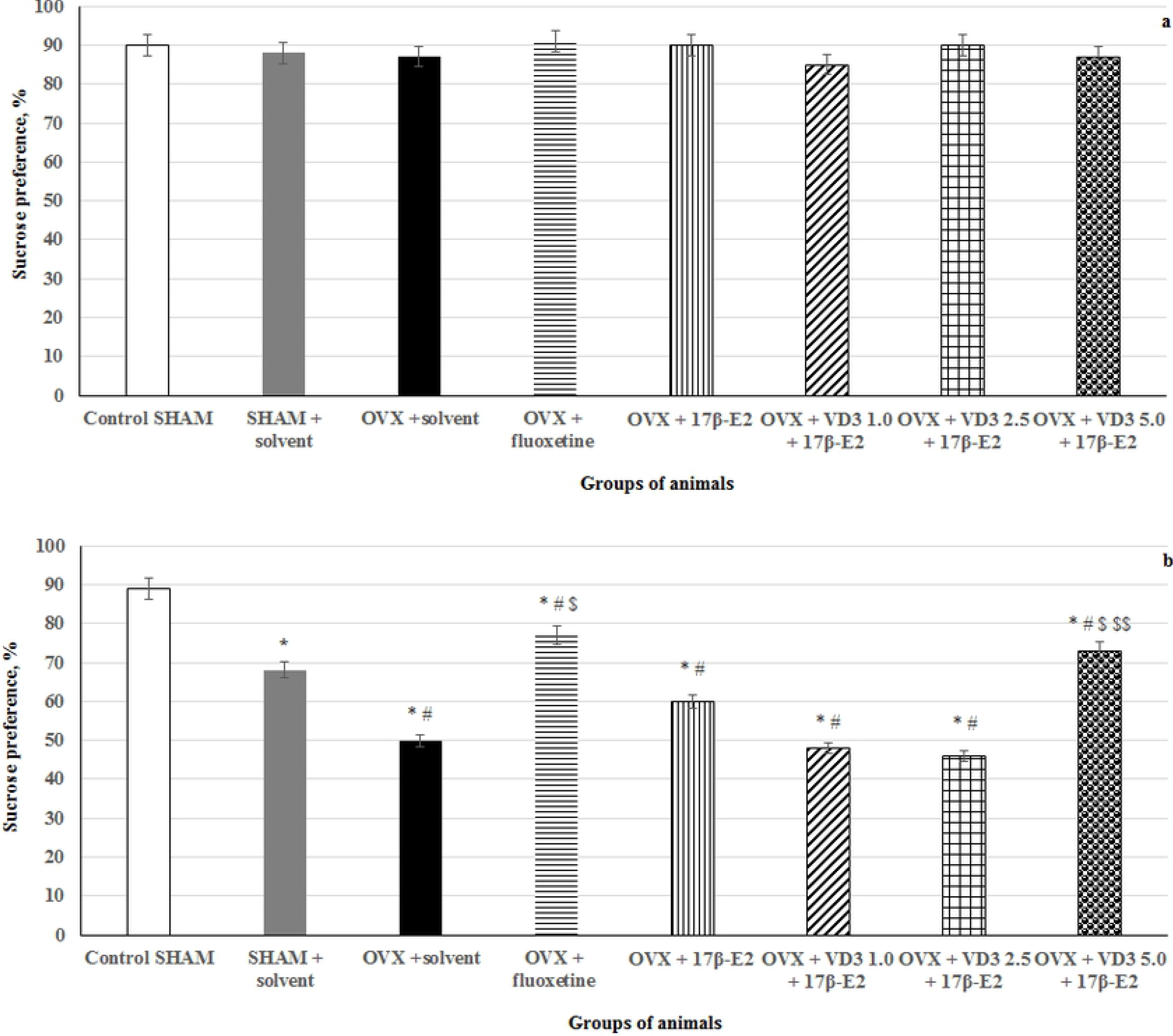
VD_3_ increases sucrose preference in the long-term OVX rats treated with 17β-E_2_ exposed to CUMS. a – prior to CUMS, b – after CUMS. * – P < 0.05 versus the control group, # – P < 0.05 versus to the SHAM group with CUMS, $ – P < 0.05 versus to the OVX group with CUMS, $$– P < 0.05 versus to the OVX group with CUMS treated with 17β-E_2_. The data are presented as mean ± SD; n = 7 in each group.

### VD_3_ decreases depression-like behavior in the forced swimming test of long-term OVX rats treated with 17β-E_2_ exposed to CUMS

CUMS produced a significant increase of the immobility time and decrease of swimming time in the long-term OVX rats compared to the non-CUMS/CUMS SHAM rats (F(1,34) = 52.84, F(1,76) = 68.89, respectively, P<0.05; Fig. 4). VD_3_ (5.0 mg/kg), as well as fluoxetine treatment, significantly reduced the immobility time and increased the swimming time in the long-term OVX treated with 17β-E_2_ compared to the OVX plus solvent or 17β-E_2_ /SHAM with CUMS groups (P < 0.05; Fig. 4). We did not find any effects of VD_3_ (1.0 or 2.5 mg/kg) administration on the depression-like parameters of the long-term OVX rats exposed to CUMS treated with low dose of 17β-E_2_ in the FST compared to the OVX plus solvent or 17β-E_2_ /SHAM with CUMS groups (P > 0.05; Fig. 4). There was no difference in the climbing time in all the experimental groups compared to the OVX/SHAM with CUMS groups (P > 0.05; Fig. 4).

**Fig. 4.**
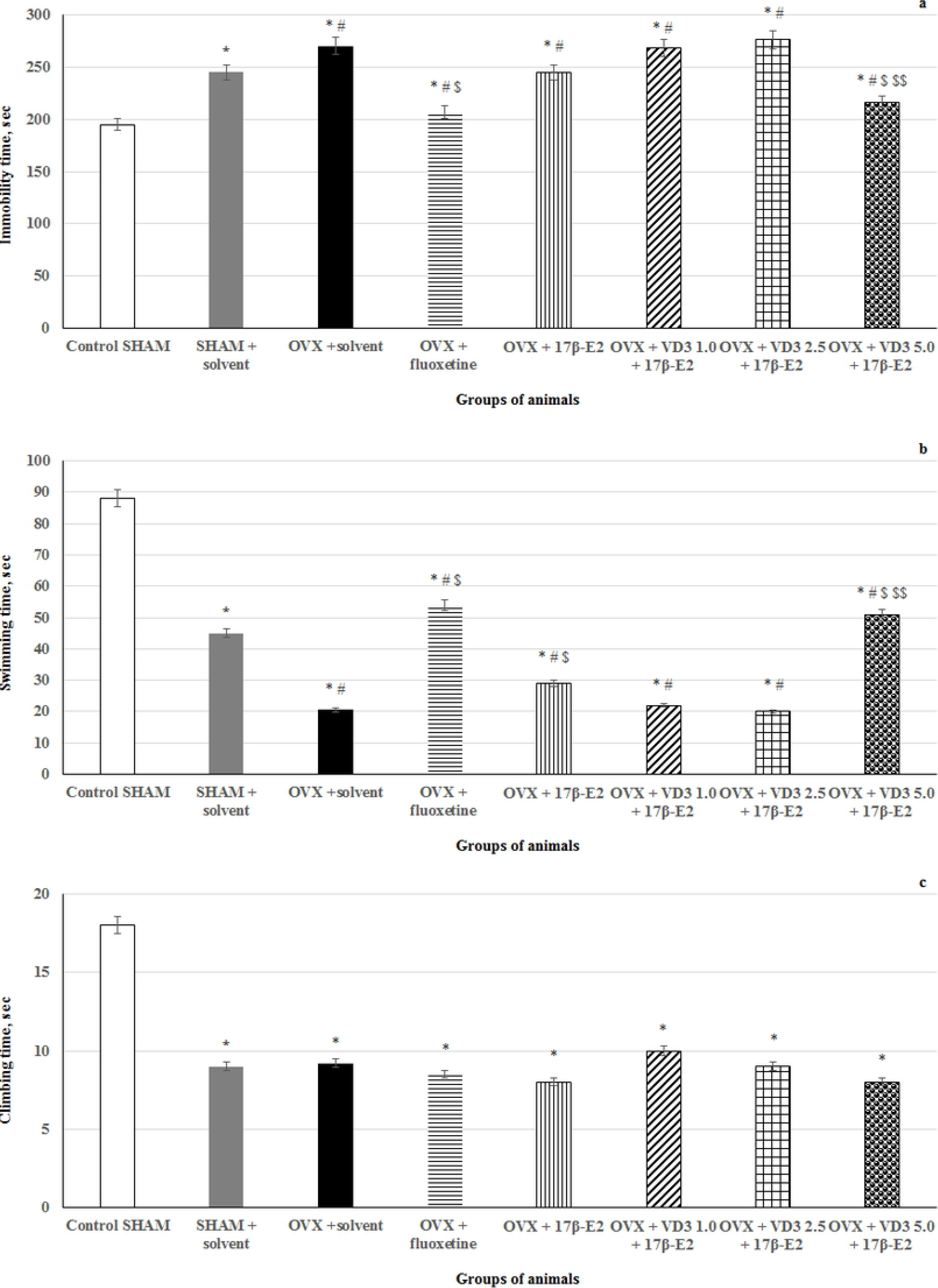
VD_3_ decreased depression-like behavior in the forced swimming test of long-term OVX rats treated with 17β-E_2_ exposed to CUMS. a – immobility time, sec, b – swimming time, c – climbing time, sec. * – P < 0.05 versus the control group, # – P < 0.05 versus to the SHAM group with CUMS, $ – P < 0.05 versus to the OVX group with CUMS, $$– P < 0.05 versus to the OVX group with CUMS treated with 17β-E_2_. The data are presented as mean ± SD; n = 7 in each group.

### VD_3_ changes the behavior in the open field test of long-term OVX rats treated with 17β-E_2_ exposed to CUMS

Following 28 days of CUMS protocol, there were no statistically significant differences for grooming activities between all the experimental groups of rats in the OFT (F(1,34) = 0.82, P > 0.05; Fig. 5). The long-term OVX rats with CUMS showed a decreased number of rearings and crossings when they were compared to the non-CUMS/CUMS SHAM groups (F(1,34) = 14.14, P < 0.05; Fig. 5). Administration of fluoxetine, 17β-E_2_, as well as treatment with VD_3_ at various doses significantly elevated the number of rearings and crossings in the long-term OVX rats with CUMS compared to the OVX/SHAM rats with CUMS plus solvent (Fig. 5).

**Fig 5.**
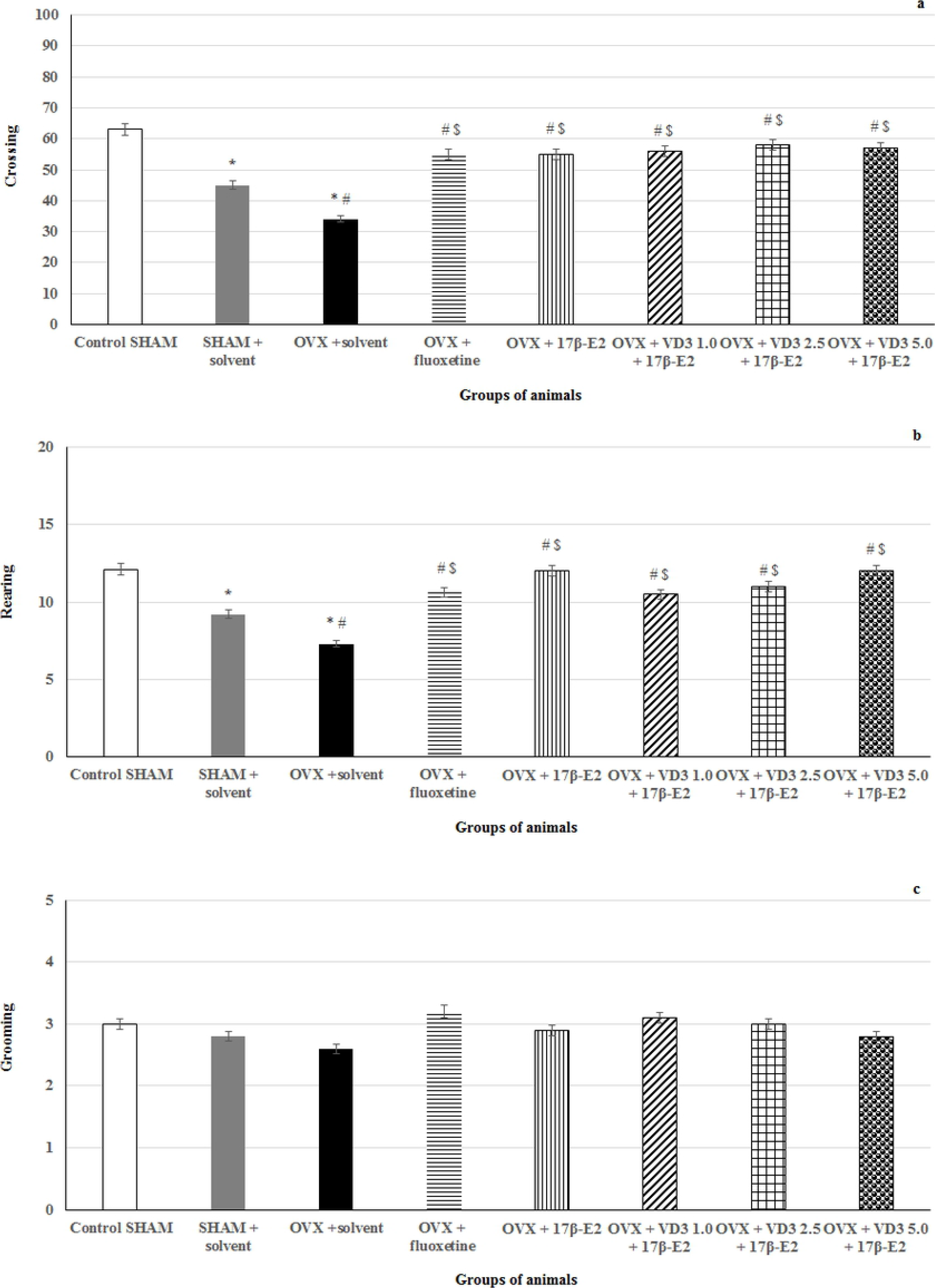
VD_3_ alters the behavior in the open field test of long-term OVX rats treated with 17β-E_2_ exposed to CUMS. a – crossing, b – rearing, c – grooming. * – P < 0.05 versus the control group, # – P < 0.05 versus to the SHAM group with CUMS, $ – P < 0.05 versus to the OVX group with CUMS, $$– P < 0.05 versus to the OVX group with CUMS treated with 17β-E_2_. The data are presented as mean ± SD; n = 7 in each group.

### VD_3_ alters serum corticosterone and VD_3_ levels in long-term OVX rats treated with 17β-E_2_ exposed to CUMS

The ELISA assay revealed that serum CS levels were elevated, it also indicated decreased VD_3_ concentrations in the long-term OVX rats with CUMS compared to the non-CUMS/CUMS SHAM groups (F(1,34) = 78.56, F(1,34) = 56.12, respectively, P < 0.05; Fig. 6a). VD_3_ did however not effect upon the pathologically elevated CS levels in the blood serum among the long-term OVX females exposed to CUMS treated with 17β-E_2_ when these were compared to the OVX plus solvent or 17β-E_2_ /SHAM with CUMS rats (P < 0.05 Fig. 6b). Fluoxetine did not change the serum VD_3_ levels, but significantly reduced the serum CS levels in the long-term OVX rats plus 17β-E_2_ exposed to CUMS compared to the OVX plus solvent or 17β-E_2_ /SHAM with CUMS rats (P < 0.05; Fig. 6).

**Fig. 6.**
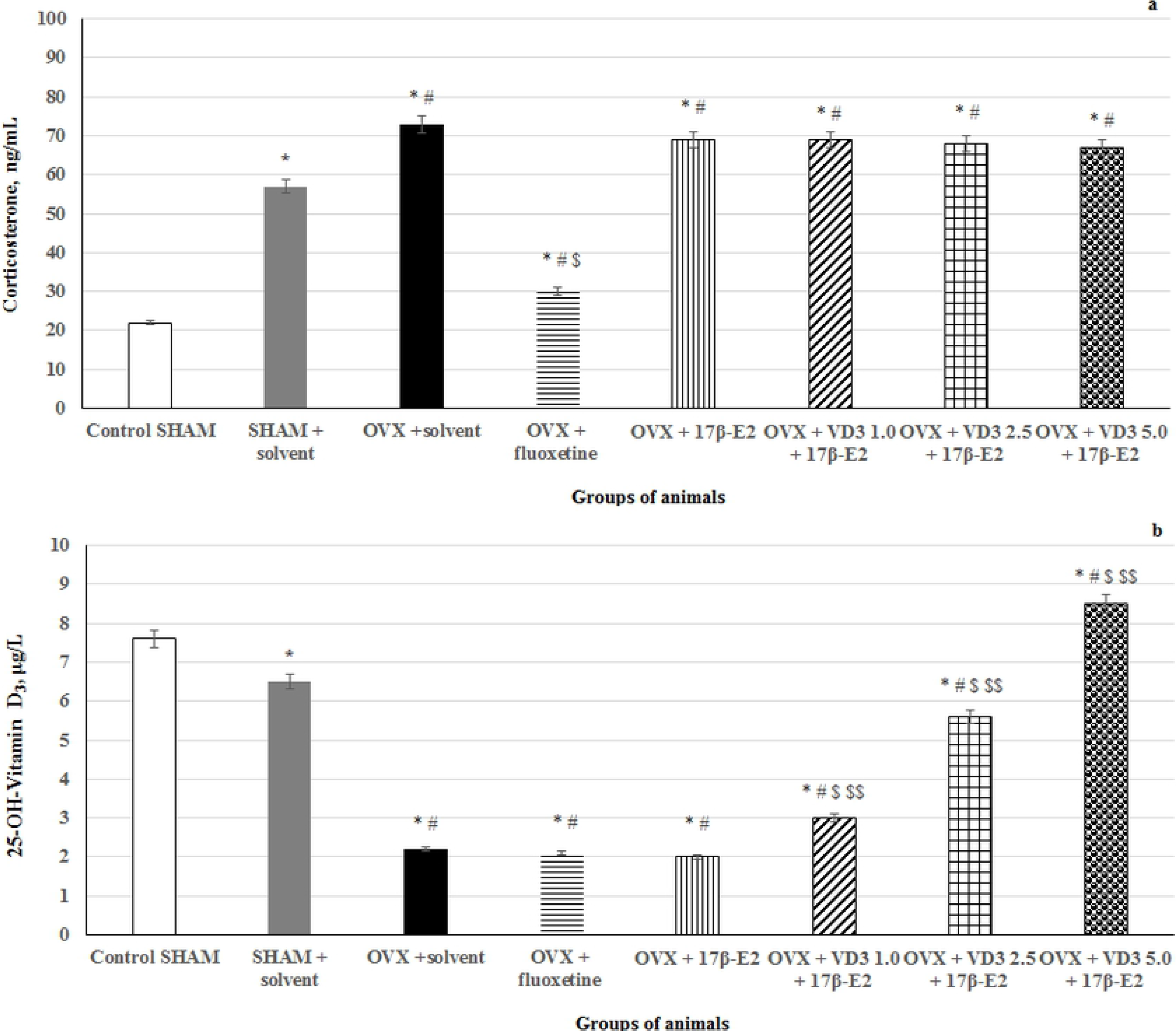
VD_3_ alters serum corticosterone and VD_3_ levels in long-term OVX rats treated with 17β-E_2_ exposed to CUMS. a – corticosterone, ng/ml, b – 25-OH-VD3, μg/ml. * – P < 0.05 versus the control group, # – P < 0.05 versus to the SHAM group with CUMS, $ – P < 0.05 versus to the OVX group with CUMS, $$– P < 0.05 versus to the OVX group with CUMS treated with 17β-E_2_. The data are presented as mean ± SD; n = 7 in each group.

### VD_3_ modulates hippocampal BDNF levels/protein expression in long-term OVX rats treated with 17β-E_2_ exposed to CUMS

CUMS significantly reduced BDNF concentrations in the hippocampus of SHAM rats compared to the non-CUMS control females (P < 0.05; Fig. 7). CUMS produced a decrease of hippocampal BDNF levels in the long-term OVX rats compared to the non-CUMS/CUMS SHAM rats (F(1,34) = 28.44, P < 0.05; Fig. 7). Supplementation with VD_3_ (5.0 mg/kg) or fluoxetine (10.0 mg/kg) increased BDNF levels in the hippocampus of long-term OVX rats treated with low dose of 17β-E_2_ exposed to CUMS compared to the OVX plus solvent or 17β-E_2_/SHAM rats with CUMS (P < 0.05; Fig. 7). No signicant differences of VD_3_ (1.0 and 2.5 mg/kg) were found on hippocampal BDNF levels of the long-term OVX rats treated with 17β-E_2_ exposed to CUMS compared to the long-term OVX with CUMS rats plus solvent (P > 0.05; Fig. 7).

**Fig. 7.**
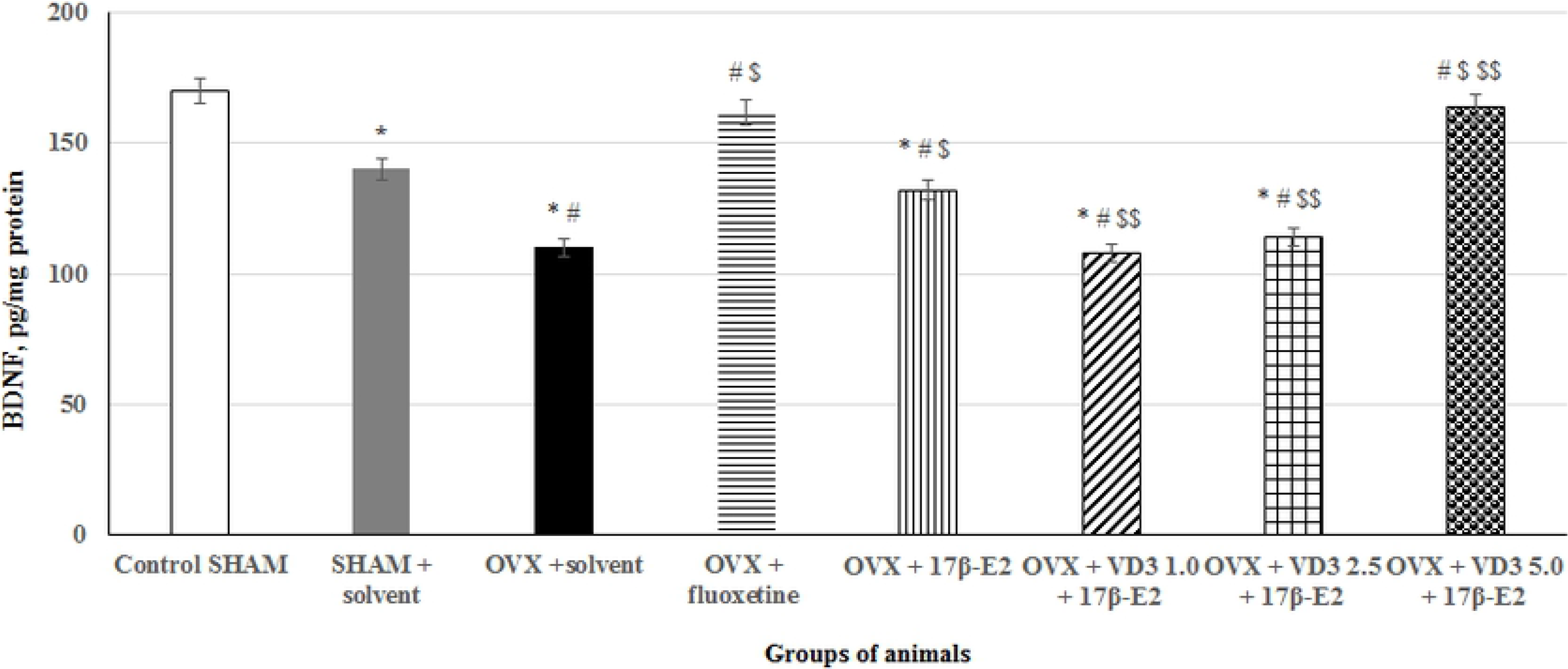
VD_3_ modulates hippocampal BDNF levels in long-term OVX rats treated with 17β-E_2_ exposed to CUMS tested by ELISA. * – P < 0.05 versus the control group, # – P < 0.05 versus to the SHAM group with CUMS, $ – P < 0.05 versus to the OVX group with CUMS, $$– P < 0.05 versus to the OVX group with CUMS treated with 17β-E_2_. The data are presented as mean ± SD; n = 7 in each group.

Western blotting analysis revealed that BDNF protein levels in the hippocampus of SHAM rats submitted to CUMS were lower compared to non-CUMS control females (P < 0.05; Fig. 8). BDNF protein levels were decreased in the hippocampus of long-term OVX rats with CUMS compared to the non-CUMS/CUMS SHAM rats (F(1,34) = 34.45, P<0.05; Fig. 8). VD_3_ (5.0 mg/kg) or fluoxetine (10.0 mg/kg) resulted in significant elevated levels of hippocampal BDNF protein expression in long-term OVX plus 17β-E_2_ rats compared to the OVX plus solvent or 17β-E_2_/SHAM rats with CUMS (P < 0.05; Fig. 8). There were no signicant differences of VD_3_ (1.0 and 2.5 mg/kg) supplementation on the BDNF protein levels in the hippocampus of the long-term OVX rats treated with 17β-E_2_ exposed to CUMS compared to the long-term OVX with CUMS rats plus solvent (P > 0.05; Fig. 8).

**Fig. 8.**
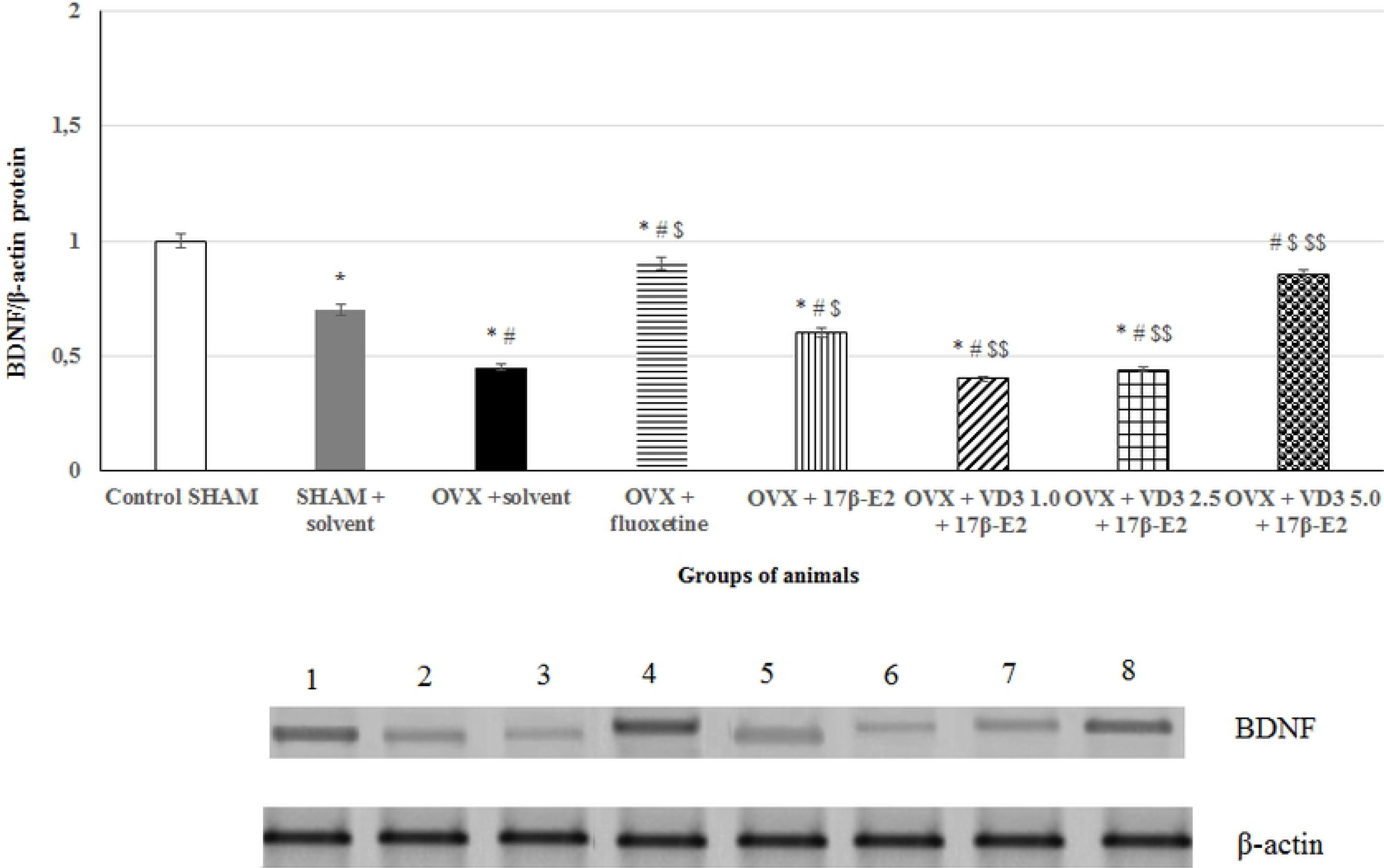
VD_3_ modulates hippocampal BDNF expression in long-term OVX rats treated with 17β-E_2_ exposed to CUMS detected with western blotting. 1 – control SHAM, 2 – SHAM + CUMS + solvent, 3 – OVX + CUMS + solvent, 4 – OVX rats + CUMS + fluoxetine, 5 – OVX rats + CUMS + 17β-E_2_, 6 – OVX rats + CUMS + VD_3_ 1.0 mg/kg + 17β-E_2_, 7 – OVX rats + CUMS + VD_3_ 2.5 mg/kg + 17β-E_2_, 8 – OVX rats + CUMS + VD_3_ 5.0 mg/kg + 17β-E_2_. * – P < 0.05 versus the control group, # – P < 0.05 versus to the SHAM group with CUMS, $ – P < 0.05 versus to the OVX group with CUMS, $$– P < 0.05 versus to the OVX group with CUMS treated with 17β-E_2_. The data are presented as mean ± SD; n = 7 in each group.

### VD_3_ modulates hippocampal 5-HT and 5-HIAA levels in long-term OVX rats treated with 17β-E_2_ exposed to CUMS

HPLC assay showed that CUMS decreased 5-HT levels in the hippocampus of SHAM rats compared to non-CUMS control females (P < 0.001; Fig. 9a). In addition, the SHAM rats with CUMS showed a significant enhanced 5-HT turnover rate in the hippocampus compared to the non-CUMS control group. The post-hoc test found that long-term OVX rats with CUMS showed a more markedly decrease of 5-HT concentrations in the hippocampus compared to the non-CUMS/CUMS SHAM rats (F(1,34) = 22.84, P<0.001; Fig. 9a). A significant increase of 5-HIAA contents were detected in the hippocampus of long-term OVX rats with CUMS rats compared to the non-CUMS/CUMS SHAM rats (F(1,34) = 34.56, P<0.001; Fig. 9b). VD_3_ (5.0 mg/kg) or fluoxetine (10.0 mg/kg) resulted in normalization of 5-HT and 5-HIAA levels in long-term OVX rats plus low dose of 17β-E_2_ compared to the OVX plus solvent or 17β-E_2_/SHAM rats with CUMS (P < 0.001; Fig. 9). Neither VD_3_ (1.0 mg/kg) nor VD_3_ (2.5 mg/kg) altered 5-HT and 5-HIAA levels of the long-term OVX rats given with 17β-E_2_ exposed to CUMS when these values were compared to similar parameters of the long-term OVX rats with CUMS plus solvent (P > 0.001; Fig. 9).

**Fig. 9.**
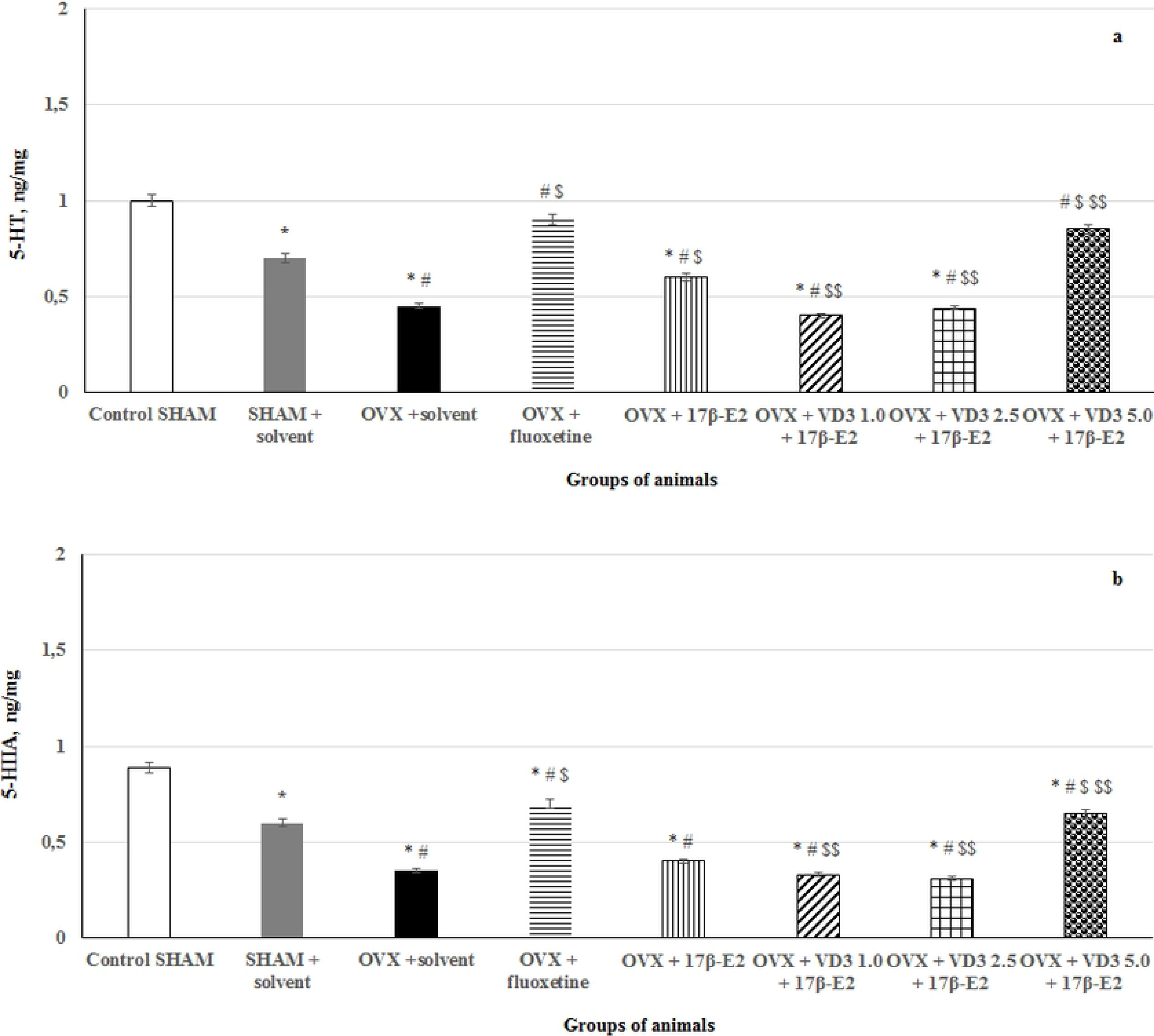
VD_3_ modulates hippocampal 5-HT and 5-HIIA levels in long-term OVX rats treated with 17β-E_2_ exposed to CUMS tested by HPLC. a – 5-HT, ng/mg wet tissue, b – 5-HIIA, ng/mg wet tissue. * – P < 0.05 versus the control group, # – P < 0.05 versus to the SHAM group with CUMS, $ – P < 0.05 versus to the OVX group with CUMS, $$– P < 0.05 versus to the OVX group with CUMS treated with 17β-E_2_. The data are presented as mean ± SD; n = 7 in each group.

The present preclinical study analyzed the antidepressant-like effects of VD_3_ treatment (1.0, 2.5 and 5.0 mg/kg, s.c.) in long-term adult OVX female rats with low dose of 17β-E_2_ subjected to the CUMS. A CUMS paradigm is a well-known experimental paradigm that imposes pathophysiological impairments in mood state, which are relevant for the understanding of clinical depressive disorders in humans [47–49]. In the present work, the implications of BDNF signaling pathway, as well as the 5-HT system, in the mechanisms of VD_3_ activity in depression were tested with regards to the affective-related condition of long-term adult OVX rats who were administered a low dose of 17β-E_2_ and exposed to CUMS.

The results of this study showed that there were marked anhedonia-/depression-like behaviors in the adult long-term OVX rats undergoing CUMS, as assessed by SPT and LDT, respectively. Moreover, long-term OVX rats undergoing CUMS exhibited decreased locomotor and rearing activities in the OFT. The ELISA assay clearly demonstrated elevated serum CS levels, as well as lower VD_3_ concentrations in adult long-term OVX rats subjected to CUMS. In addition, a decreased BDNF concentration/protein expression and a decreased 5-HT turnover were found in the hippocampus of the long-term OVX rats exposed to CUMS. The results of this study confirm that CUMS produces marked behavioral, neuroendocrine, and neurochemical changes in adult OVX rats with long-term ovarian hormone deficiency (post-ovariectomy period of 3 months). These findings are in agreement with other studies concerning effects of long-term estrogen deprivation in female rodents subjected to a CUMS procedure results in a profound affective-like profile [57,58].

Administration of 17β-E_2_ failed to completely restore behavioral and biochemical parameters in the long-term OVX rats exposed to CUMS. Fluoxetine decreased anhedonia-like and depression-like states, increased BDNF/5-HT and reduced CS levels in the hippocampus of the long-term OVX female rats exposed to CUMS. Other studies have demonstrated that fluoxetine reversed depression-like profiles of OVX rats in a stress depression model [52]. Moreover, preclinical findings have demonstrated that fluoxetine might restore the functional activity of the HPA and 5-HT systems, and BDNF expression in different structures of the brain, improving symptoms of depression in OVX rats with different postovariectomy intervals in both stressed and non-stressed behavioral models [52,53,58]. The results of the current study indicate that CUMS provokes marked behavioral, and neurochemical/neurohormonal alterations in adult OVX rats with long-lasting estrogens decline. Our data are in agreement with our recent data and findings of other studies which indicated that long-term estrogen deprivation in female rodents subjected to a CUMS procedure results in a profound affective-like profile [57,58].

The most important conclusion that can be made based on the present study are that VD_3_ treatment has antidepressant-like effects in long-term adult OVX rats treated with low dose of 17β-E_2_ subjected to CUMS. VD_3_ given with a dose of 5.0 mg/kg reversed anhedonia-like and depression-like states in the SPT/LDT paradigms in the long-term OVX rats treated with 17β-E_2_ subjected to CUMS, which were similar to the effects of the fluoxetine treatment. Moreover, the VD_3_ application reversed the behavioral impairments observed in the OFT in the long-term OVX rats supplemented with 17β-E_2_ subjected to CUMS. Neurochemical and biochemical assays found that VD_3_ (5.0 mg/kg, s.c.) increased the serum VD_3_ levels and hippocampal 5-HT/BDNF levels, but did not modify the serum CS content in the long-term OVX rats treated with 17β-E_2_ and exposed to CUMS. Western blot analysis revealed that VD_3_ increased BDNF protein expression in the hippocampus of long-term OVX rats treated with 17β-E_2_ subjected to CUMS. These data suggest that VD_3_ attenuates the CUMS-produced behavioral impairments and normalized the serum VD_3_ levels, as well as 5-HT and BDNF production in the hippocampus of long-term OVX rats treated with 17β-E_2_ and exposed to CUMS.

The neurochemical and behavioral effects of VD_3_ at doses of 1.0 or 2.5 mg/kg were profoundly different to the effects of VD_3_ at a dose of 5.0 mg/kg. VD_3_ at these two doses failed to change the neurochemical and behavioral parameters of the long-term OVX rats administered with low dose of 17β-E_2_ who were exposed to a CUMS procedure. The fact that the VD_3_ application in all tested doses increased the rearing and crossing allows us to conclude that the effects of VD_3_ in the SPT and EPM/LDT cannot possibly be attributed to behavioral changes in the OFT, but rather should be interpreted as a direct expression of the anhedonia and depression-like behavior in the long-term OVX rats treated with low dose of 17β-E_2_ and exposed to CUMS. Interestingly, the results of behavioral and neurochemical effects of VD_3_ at dose of 1.0 mg/kg in the long-term OVX rats treated with low dose of 17β-E_2_ and exposed to CUMS are yielding an opposite pattern from the results of our recent study on the long-term OVX rats with CUMS but without hormonal supplementation. More specifically, VD_3_ at dose of 1.0 mg/kg produced more markedly depression-like levels in the long-term OVX rats submitted to CUMS without hormonal treatment, but it did not induce any aggravation of depressive symptoms in the OVX rats plus CUMS who were given 17β-E_2._ However, activity of VD_3_ at doses of 2.5 and 5.0 mg/kg on depression-like behavior were alike the effects that VD_3_ has for the long-term OVX rats subjected to CUMS treated with solvent or low dose of 17β-E_2_.

Moreover, unexpectedly, the effects of VD_3_ administered at a dose of 5.0 mg/kg on the serum CS levels were contrasting in the long-term OVX rats plus CUMS treated with hormonal preparation or solvent. Although, in the current study, VD_3_ at a dose of 5.0 mg/kg improved behavioral and neurochemical states of the long-term OVX rats treated with a low dose of 17β-E_2_, we did not observe any changes of the serum CS levels when this dose was applied to animals.

This is the first study to compare the role of VD_3_ on the behavioral and neurochemical consequences of a CUMS procedure in adult long-term OVX rats treated with 17β-E_2_ and exposed to a CUMS procedure. Further studies are strongly recommended to explore why the behavioral effects of VD_3_ supplementation at different doses produce very different effects in the long-term OVX rats submitted to CUMS at the presence or absence of hormonal treatment. On the other hand, our earlier study demonstrated that VD_3_ treatment decreased depression-like behavior in the long-term, non-stressed OVX rats [42,43]. Thus, the data of the present study when using long-term OVX rats subjected to CUMS are contradictory to the findings of our previous study concerning the antidepressant-like effects of VD_3_ at similar doses in non-stressed long-term OVX rats [43]. Together, the results of the present study and our previous work indicate that the behavioral effects of VD_3_ are dependent on used experimental paradigms (stressed or non-stressed) and hormonal treatment (absence or presence). This fact (concerning the various effects of VD_3_ on the depression-like state in the long-term OVX rats that has been observed in both our studies) might provide further explanation findings concerning the antidepressant-like effects of VD_3_ in the animal and human studies on female subjects with long-lasting estrogen deficiency, especially, when there exists additional hormonal treatment. Further studies are, however, required to consider why the behavioral effects of VD_3_ supplementation are different in non-stressed and stressed long-term OVX rats supplemented with hormonal treatment or not.

Based on the present results, it might be assumed that the beneficial implications of VD_3_ in the long-term OVX rats treated with low dose of 17β-E_2_ with CUMS is associated with the synergic effects of VD_3_ and 17β-E_2_ on the BDNF/5-HT pathways. Estrogens, as well as VD_3_, controls functional activity of the BDNF/5-HT systems in the structures of the brain which are involved in the neurobiological mechanisms of affective-related disorders [59–61]. Such complex effects of VD_3_ on BDNF/5-HT pathways might promote a greater effect of the combination of VD_3_ plus 17β-E_2_ than applications of only 17β-E_2_.

This is the first study to show the action of VD_3_ on the behavioral and biochemical consequences of a CUMS in adult long-term OVX rats treated with low dose of 17β-E_2_. The restoration BDNF/5-HT activity by VD_3_ treatment is a promising fact for the study of treatment of affective-related diseases that are associated with low levels of VD_3_. These results suggest that normalization of the functional activity of BDNF/5-HT pathways for VD_3_ may be one of the fundamental components of its activity. The ameliorating effect of VD_3_ on CUMS-induced impairments of the BDNF/5-HT pathways in the hippocampus of the long-term OVX rats treated with low dose of 17β-E_2_ are strongly in accordance with the monoamines and BDNF hypotheses of depression [62–64]. Since, both VD_3_ and 17β-E_2_ have anti-inflammatory activity, one another mechanisms by which VD can improve depression-like symptoms in long-term OVX rats could be by modulation of NF-kB/p65 signaling [59,61–63]. Moreover, there exists a complex interaction between BDNF and NF-kB in the cellular mechanisms of depression development [59,65–66]. It is hence possible that VD_3_ and 17β-E_2_ in the long-term OVX rats are associated with genomic cross-talk protein interactions via NF-kB.

To our knowledge this is the first laboratory research which indicates synergic antidepressant-like effect of VD_3_ at dose of 5.0 mg/kg in long-term OVX rats submitted CUMS treated with 17β-E_2_. It is well-known that exposure to stressful triggers profoundly alters affective-related states, and predisposes an individual to anxiety, irritability, anhedonia, or depression [1–3]. Modern life, especially in women [3,4], is associated with numerous stressful and negative emotional situations, thereby inducing chronic exposure to stressful stimuli. It should be emphasized that affective-related deteriorations become more severe in postmenopausal women since the female gonadal hormones physiologically acts as a potent antidepressant-like factor [8–10].

Daily aversive stimuli are considered to alter the mRNA levels of 5-HT transporters and enzymes in the brain and are thereby associated to affective-related disorders [9,10]. Moreover, stress stimuli profoundly alter the expression of BDNF, which is the key factor for neurogenesis in the hippocampus [59–61]. A reduction of the BDNF expression is linked with depression in humans [31–35]. On the contrary, an increase in BDNF expression improves stress-induced depressive disorder [33–35]. Furthermore, BDNF expression is connected with 5-hydroxytryptamine receptor expression in the brain, particularly in the hippocampus [33,59–61]. Therefore, BDNF is documented to be a key neurotrophic factor that modulates a depressive mood through its activity in the hippocampus. Clinical studies suggest that serum BDNF is more decreased in patients with depression, but antidepressant therapy produces an increase in BDNF levels [35,36,59]. Therefore, VD supplementation may be very useful for the treatment of affect-related disorders among postmenopausal females with VD deficiencies or insufficiencies and supplemented with MHT. However, more research on this topic is still required as the exact role of VD in affect related disorders associated with hormonal changes in women has not been completely understood.

## Conclusions

This study demonstrated that VD_3_ at high concentration in a combination with low dose of 17β-E_2_ had a synergic antihedonic- and antidepressant-like effects in the adult female rats following long-term ovariectomy submitted to CUMS. In contrast to fluoxetine action on hormonal state, treatment with VD_3_ at all investigated doses did not alter CS levels in the serum of long-term OVX rats exposed to CUMS treated with low dose of 17β-E_2_. Although, VD_3_ at high concentration in a combination with low dose of 17β-E_2_ restored BDNF levels and 5-HT/5-HIAA contents, as well as 5-HT turnover in the hippocampus of long-term OVX rats exposed to CUMS treated. Thus, this is the first study in long-term OVX female rats showing the completely opposite effects of VD_3_ on depression-like behavior that are depended used dose and presence/absence of stressful factors. Our study yields new knowledge into the mechanisms by which VD_3_ affects to alleviate anhedonia- and depressive-like behaviors in female rodents with long-term estrogen deficiency in stress model of depression, with a specific effects on behavioral state via 5-HT and BDNF pathways in the hippocampus. In animal model of depression produced by chronic mild stress, VD_3_ may not change CS levels, however, may produce effects on depression-like behavior that result from 5-HT neurotransmitter brain system activation and BDNF signaling, through mechanisms which distinguish from those of classical selective serotonine reuptake inhibitors (SSRIs) like fluoxetine.

## Data Availability Statement

The protocols of behavioral and biochemical measurements used to support the findings of this study are available from the corresponding author upon request.

